# Disrupted Neurovisceral Integration and Emotional Processing in Early Cerebral Small Vessel Disease

**DOI:** 10.1101/2025.10.03.678529

**Authors:** Olga Dobrushina, Larisa A. Dobrynina, Galina A. Arina, Viktoriya V. Trubitsyna, Evgenia S. Novikova, Vlada V. Kolomoitseva, Daria A. Alexandrova, Mariia V. Gubanova, Elena I. Kremneva, Marina V. Krotenkova

**Author notes:** Correspondence: Olga Dobrushina, 40 George Street, Glasgow, UK, G1 1QE.

## Abstract

**Background:** Cerebral small vessel disease (cSVD) is a prevalent age-related microangiopathy leading to cognitive decline. Beyond established mechanisms and risk factors, disturbances in neurovisceral integration—the brain–body coordination of emotional and cardiovascular responses—may represent a systems-level pathway relevant to cSVD pathogenesis. We therefore investigated whether early cSVD is characterised by altered neurovisceral integration and emotional processing.

**Methods:** Eighty-two adults were studied: middle-aged adults with early cSVD (n=37), age-matched adults without brain abnormalities (n=23), and young healthy adults (n=22). Participants completed an ecologically grounded social emotion fMRI paradigm alongside self-report measures of interoception and emotional processing. Neurovisceral integration was operationalised as arousal-related modulation of right anterior insula activity and concurrent heart rate dynamics during the task.

**Results:** Early cSVD, but not healthy ageing, was associated with diminished insular encoding of arousal, altered heart-rate dynamics, reduced emotional differentiation, and loss of the age-related positivity effect. Self-report questionnaires showed parallel alterations in everyday functioning. Effects remained significant after adjustment for cardiovascular risk factors and cognitive performance.

**Conclusion:** Study findings indicate disruptions in neurovisceral integration and emotional processing across neural, physiological, and behavioural levels, identifying them as candidate biomarkers of early cSVD and potential targets for preventive interventions.

## Introduction

Cerebral small vessel disease (cSVD) is a common age-related microangiopathy characterised by white matter injury and disruption of fronto-subcortical networks.^1^ The prevalence of cSVD increases with age, reaching nearly 100% by age 90, making it a major contributor to cognitive decline and dementia in later life.^2^ While cognitive impairment has been the primary focus of cSVD research, affective disturbances are increasingly recognised as clinically relevant features of the disorder. Beyond cognitive deficits, cSVD may involve disruption of large-scale brain networks supporting emotional and autonomic regulation^3^ and is commonly associated with depression.^4, 5^

Of relevance to vascular mechanisms underlying cSVD, major depressive disorder has been consistently recognised as a risk factor for hypertension, coronary heart disease, and stroke.^6^ Psychotherapeutic treatment of depression has resulted in reduced cardiovascular risk, suggesting that emotional disturbances may have a causal role in vascular pathology.^7^ Difficulties in emotional awareness, such as alexithymia, have likewise been identified as independent predictors of arterial hypertension and cardiovascular mortality.^8, 9^ Together, these findings suggest that in cSVD, affective alterations may not only follow brain injury but also contribute to disease progression through cardiovascular dysregulation, while effective emotional processing may represent a protective factor.

In healthy ageing, emotional processing is characterised by increased differentiation of emotional experience and a shift toward more positive affect. Throughout adulthood, emotion regulation becomes more context-sensitive and effective, leading to differentiated, adaptive use of strategies.^10^ With a shift in motivation towards meaningful experiences, the overall emotional tone changes to more positive, a phenomenon known as the age-related positivity effect.^11^ These findings of improved self-regulation are consistent with epidemiological evidence showing a decline in the prevalence of depression and anxiety from early to late adulthood.^12^ Affective changes in cSVD may therefore represent a disruption to this healthy ageing trajectory, with parallel alterations in emotional and cardiovascular regulation.

On the neural level, emotional and physiological processes are linked through neurovisceral integration. Neurovisceral integration refers to the coordinated regulation of autonomic, attentional, and affective processes by a hierarchical network involving the cortex, subcortical structures, and brainstem structures.^13, 14^ Related to this framework is interoception—the sensing and representation of internal bodily states.^15^ Distributed neural circuits integrate interoceptive signals with emotional and contextual information, enabling proactive, flexible, and differentiated regulation of physiological responses.^16, 17^ Dysfunction of this complex system has been associated with reduced autonomic adaptability, and age-related changes in these networks have been described, supporting relevance to cerebrovascular disorders.^13, 18^

Taken together, these lines of evidence suggest that cSVD pathomechanisms may involve disruptions in the physiological systems supporting neurovisceral integration and emotional processing. Such affective alterations may represent potentially modifiable mechanisms that complement established models emphasising endothelial dysfunction, inflammation, and haemodynamic factors.^1, 19^ Our previous pilot study showed a correlation between emotional intelligence and white matter integrity in middle-aged adults.^20^ Here, we investigated neurovisceral integration during emotional processing using an ecologically grounded social emotion fMRI paradigm, combined with self-report measures of everyday functioning. Emotional differentiation, the age-related positivity effect, insula-related modulation of physiological arousal, and heart rate dynamics were evaluated as candidate markers of regulatory integrity. By examining early cSVD and controlling for cognitive performance, we sought to identify affective alterations that precede cognitive impairment and may have an independent role in microvascular brain pathology.

## Material and Methods

### Study Design

A cross-sectional study included three groups of adults: middle-aged adults with early small vessel disease (middle-aged SVD+), middle-aged adults with no brain abnormalities (middle-aged SVD−), and young adults with no brain abnormalities (young SVD−). Within two weeks, participants underwent clinical evaluation, structural neuroimaging, functional magnetic resonance imaging (fMRI) with the social emotion task, and cognitive assessment.

The sample size was established to enable whole-brain analyses of continuous variables across the entire sample and between-group comparisons using region-of-interest (ROI) data. Since the fMRI task was novel, the target sample size was guided by recommendations from the neuroimaging literature, which suggests that 60–80 participants are needed to detect moderate effects in whole-brain analyses, and that ROI analyses can be adequately powered with smaller samples of 20–25 participants.^21, 22^

The study protocol was approved by the Ethics Committee and the Institutional Review Board of the Russian Center of Neurology and Neurosciences (protocols 4-6/19, 1-3/20, and 4-3/21), and all participants gave written informed consent. All procedures contributing to this work complied with the ethical standards of the relevant national and institutional committees on human experimentation and with the Helsinki Declaration of 1975, as revised in 2008. The study was not pre-registered. The data supporting the article are available from the corresponding author on request.

### Participants

#### Inclusion criteria

Three groups were defined by age (middle-aged vs. young) and cSVD status (present vs. absent): middle-aged SVD+ (40–65 years with early cSVD), middle-aged SVD– (40–65 years), and young SVD– (18–39 years).

Small vessel disease status was assessed using T2 (0.68×0.68×3 mm), sagittal 3D-T2 FLAIR (1×1×1 mm), and sagittal T1 MPRAGE (1×1×1 mm) MRI sequences, evaluated by a radiologist. Early cSVD was defined as a modified Fazekas score of 1 or 2 and intact functional independence.^23^ The Fazekas score reflects periventricular and deep white matter lesion severity: 0 – absent, 1 – small/focal, 2 – larger with some bridging, 3 – confluent or ≥20 mm lesions.^24^

#### Exclusion criteria

included a history of cardiovascular events, such as stroke or myocardial infarction, cancer or systemic diseases, endocrine diseases other than mild hypothyroidism or type 2 diabetes managed without insulin, dementia, Fazekas score 3, and structural brain abnormalities apart from cSVD, including signs of neurodegenerative diseases.

#### Recruitment

Participants were recruited in Moscow, Russia, between 2020 and 2022. Middle-aged SVD+ adults were sourced through the Russian Center of Neurology and Neurosciences database of patients with cerebrovascular conditions and social networks. Young and middle-aged SVD− adults were enrolled from the community via social networks.

### Clinical Evaluation

At enrolment, study participants were evaluated by a neurologist to collect medical history, including the absence of exclusion criteria and the presence or absence of arterial hypertension. In addition, middle-aged adults underwent 24-hour ambulatory blood pressure monitoring. Arterial hypertension was defined as the existence of a previously established diagnosis, antihypertensive medication intake, or mean systolic 24-hour ambulatory blood pressure ≥130 mmHg, or mean diastolic 24-hour ambulatory blood pressure ≥80 mmHg, in accordance with the ESC/ESH guidelines.^25^

Participants completed the Toronto Alexithymia Scale (TAS-20),^26, 27^ Multidimensional Assessment of Interoceptive Awareness (MAIA-2) ^28^, Beck Depression Inventory (BDI),^29, 30^ and the State Trait Anxiety Inventory.^31^ Cognitive assessment included the Montreal Cognitive Assessment, Rey-Osterrieth Complex Figure Test, Luria’s Memory Word Test, Verbal Fluency Test (number of words produced in 1 minute for words beginning with “S”, “B”, and animal names), Stroop Test, Trail Making Test, and Tower of London Test.^32-35^ Questionnaires and tests were administered in Russian, using previously adapted versions.

### Neuroimaging Data Acquisition and Preprocessing

#### Social emotion task

Participants completed a social emotion fMRI task comprising alternating 24 Emotion and 12 Pixel blocks (Figure S1). Emotion blocks presented 2–5 s videos of actors expressing anger, irritation, joy, or pleasure, followed by valence and arousal ratings. Front-facing videos of actors, accompanied by expressive pseudo-speech, mimicked interpersonal interaction. Pixel blocks displayed pixelated versions of the same videos with a noise soundtrack, followed by ratings of the shade and intensity of colour. Stimuli were drawn from the <u>GEMEP</u> corpus^36^ and presented using Cogent (MATLAB) on a Philips SensaVue screen. Participants responded by pressing buttons with their dominant hand.

#### Data acquisition

Neuroimaging was performed using a Siemens MAGNETOM Prisma 3T scanner with a 20-channel coil and a Philips InVivo Functional Imaging System, located at the Russian Center of Neurology and Neurosciences (Moscow). Functional images were acquired using T2*-gradient echo imaging sequences (TR 2000 ms, TE 21 ms, voxel size 2×2×2 mm, FOV 192×110 mm). The scan lasted 864 seconds and comprised 432 volumes. A gradient echo field map was acquired for distortion correction (TR = 520 ms, TE1 = 4.92 ms, TE2 = 7.38 ms, voxel size = 2×2×2 mm). A sagittal T1 MPRAGE sequence (TR 2300 msec, TE 2.98 msec, voxel size 1×1×1 mm, FOV 256×248 mm) was used as an anatomical reference and for grey matter volume evaluation. Pulse data were recorded simultaneously from a pulse oximeter located on the participant’s non-dominant hand using the built-in Siemens Physiological Measurement Unit. Due to incomplete backup following a console malfunction, pulse data from seven participants were lost (middle-aged SVD+: 6; middle-aged SVD−: 1).

#### Data preprocessing

fMRI data were processed using <u>SPM12</u> in Matlab. A standard pipeline included slice-timing correction, calculation of the voxel displacement map, realignment and unwrapping of the functional images, co-registration of the structural and functional images, spatial normalisation into standard Montreal Neurological Institute (MNI) space, and spatial smoothing using a Gaussian kernel of 8 mm full width at half maximum. Grey matter and total intracranial volumes were calculated using the <u>CAT12</u> toolbox, which performs tissue segmentation and spatial normalisation via a unified approach.^37^

### Data Analysis

#### Analysis of Behavioural Data

Data analysis and visualisation were performed in R Project using packages ‘dplyr’, ‘tidyr’, ‘purrr’, ‘lme4’, ‘lmerTest’, ‘emmeans’, ‘Hmisc’, ‘corrplot’, ‘RcmdrMisc’, ‘MASS’, ‘ggplot2’, ‘ggpubr’, ‘ggbeeswarm’, ‘ggrain’, ‘grid’, ‘gridExtra’, ‘cowplot’, ‘ggsignif’.

*Emotional differentiation* and *positivity* scores were calculated from ratings in the social emotion task. Emotional differentiation was operationalised as the variance in absolute arousal and valence ratings across trials, reflecting the extent to which emotional experience was nuanced rather than confined to broad categories such as ‘strong’ or ‘positive’. Positivity was defined by mean valence across trials.

Group differences were assessed using ANOVA, Kruskal–Wallis, or chi-square tests for normally distributed, non-normally distributed, and categorical variables, respectively, yielding an overall p-value. Normality was assessed using the Kolmogorov-Smirnov test. Pairwise post hoc comparisons focused on the effects of ageing (young vs middle-age SVD−) and cSVD (middle-age SVD− vs SVD+), using Tukey’s HSD, Wilcoxon rank-sum, or Fisher’s exact tests as appropriate. Post hoc *p*-values were corrected using the false discovery rate method (*p*-FDR).

#### First and Second-Level fMRI Analysis

During first-level analysis, Emotion and Pixel conditions were modelled as short blocks aligned to the onset and duration of each video. Realignment parameters and physiological noise regressors were included as covariates. Activation maps were computed for the Emotion > Pixel contrast and for parametric modulation of Emotion blocks by arousal and valence ratings.

Second-level ‘Emotion > Pixel’, ‘Arousal’, and ‘Valence’ activation maps were inspected to assess task validity (see Supplement). To examine neural correlates of emotional differentiation and positivity, these variables were entered as second-level covariates in the ‘Emotion > Pixel’ contrast. Mean valence was winsorized to reduce the influence of outliers. Voxel-level threshold was set at *p* <.001, with clusters considered significant at *p*-FDR <.05. Fragmented results were additionally explored at voxel-level *p* <.005 with a minimum cluster size of 50 voxels. Analyses were restricted to the grey matter mask.

#### Neurovisceral Integration

To evaluate neurovisceral integration, ROI data for the right ventral anterior insula were extracted from the arousal individual-level parametric modulation contrasts using <u>MarsBaR</u> (Figure S3). The ROI was defined by intersecting the thresholded group-level ‘Arousal’ activation map with the right ventral anterior insula region from the Deen et al. insula parcellation atlas.^38^ This region was selected based on evidence that the right ventral anterior insula encodes arousal intensity^39^ and our previous findings linking insular activation in the heartbeat detection task to early cSVD.^20^

Pulse time series were processed using the <u>TAPAS PhysIO Toolbox</u>. For each Emotion block, heart rate was computed as the mean within a 2–12 second window following video onset. Implausible heart rate values (below 40 bpm or above 120 bpm) were excluded, followed by removal of outliers defined as values exceeding 1.5×IQR below the first quartile or above the third quartile within each participant.

To assess the association between BOLD modulation in the right anterior insula and heart rate dynamics during Emotion trials, a linear mixed model was fitted with the video onset time, insular modulation of arousal strength, and their interaction as fixed effects and heart rate as the dependent variable. To examine group differences in heart rate dynamics during the task, a model was fitted with video onset time, group, and their interaction as fixed effects. In both models, heart rate was modelled as a repeated measure across trials, with participant ID included as a random intercept.

To assess group differences in interoceptive self-report profiles, we analysed MAIA-2 subscales using multivariate analysis of variance (MANOVA) and linear discriminant analysis (LDA). Separate comparisons were performed between young adults and SVD− middle-aged adults (effect of age), and between SVD− and SVD+ middle-aged adults (effect of cSVD). MANOVA tested for multivariate group differences across eight MAIA-2 subscales. LDA was used to estimate the discriminant function and cross-validated classification accuracy, providing an interpretable profile of subscales contributing to group differences.

## Results

### Participants

The study included 82 participants: 37 middle-aged SVD+ (Fazekas 1: 32; Fazekas 2: 5), 23 middle-aged SVD−, and 22 young adults without brain abnormalities (Table 1). The sample included more females among middle-aged adults, reflecting greater willingness to participate and the demographic profile of the recruitment database. All participants identified as White.

**Table 1.**
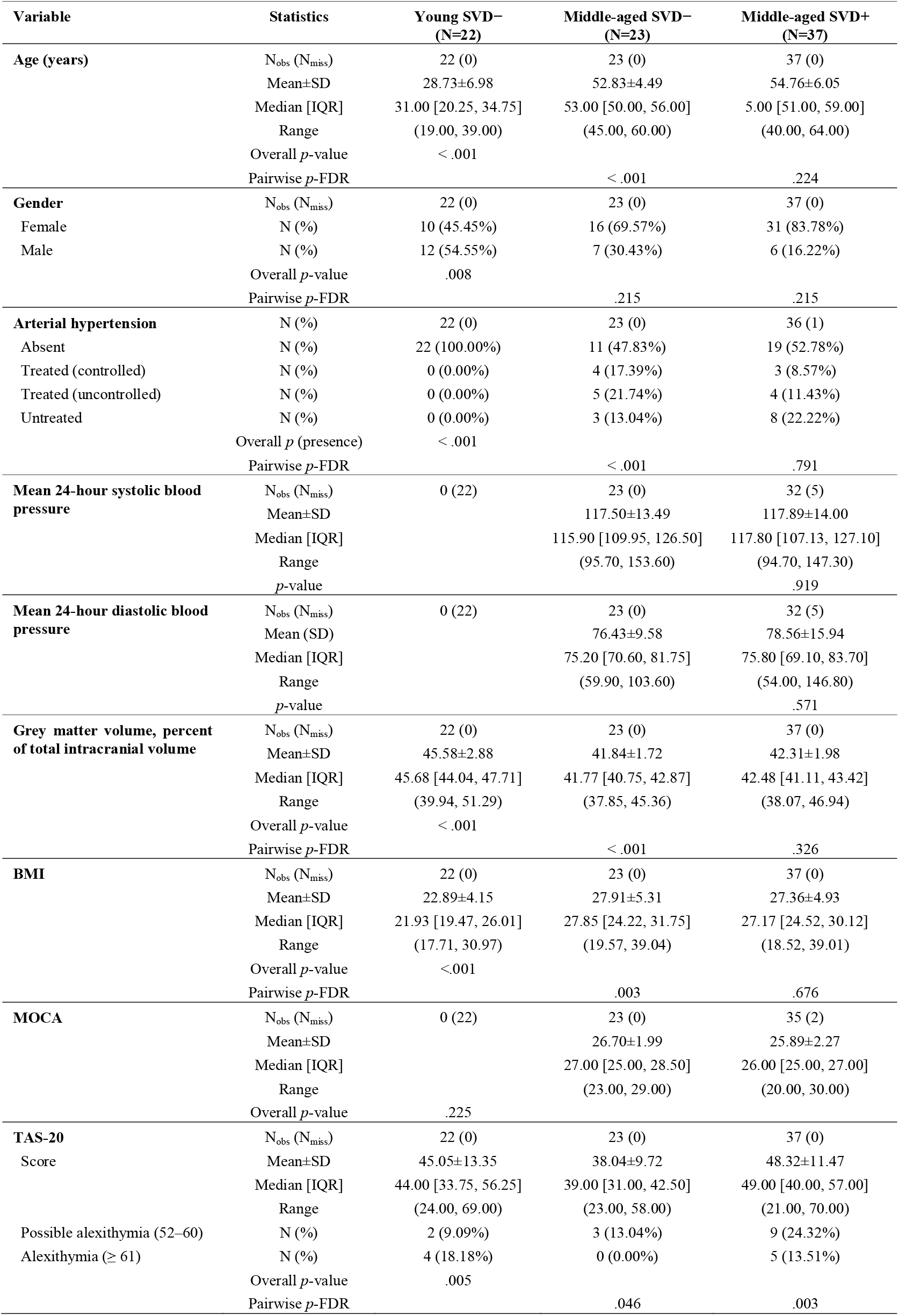
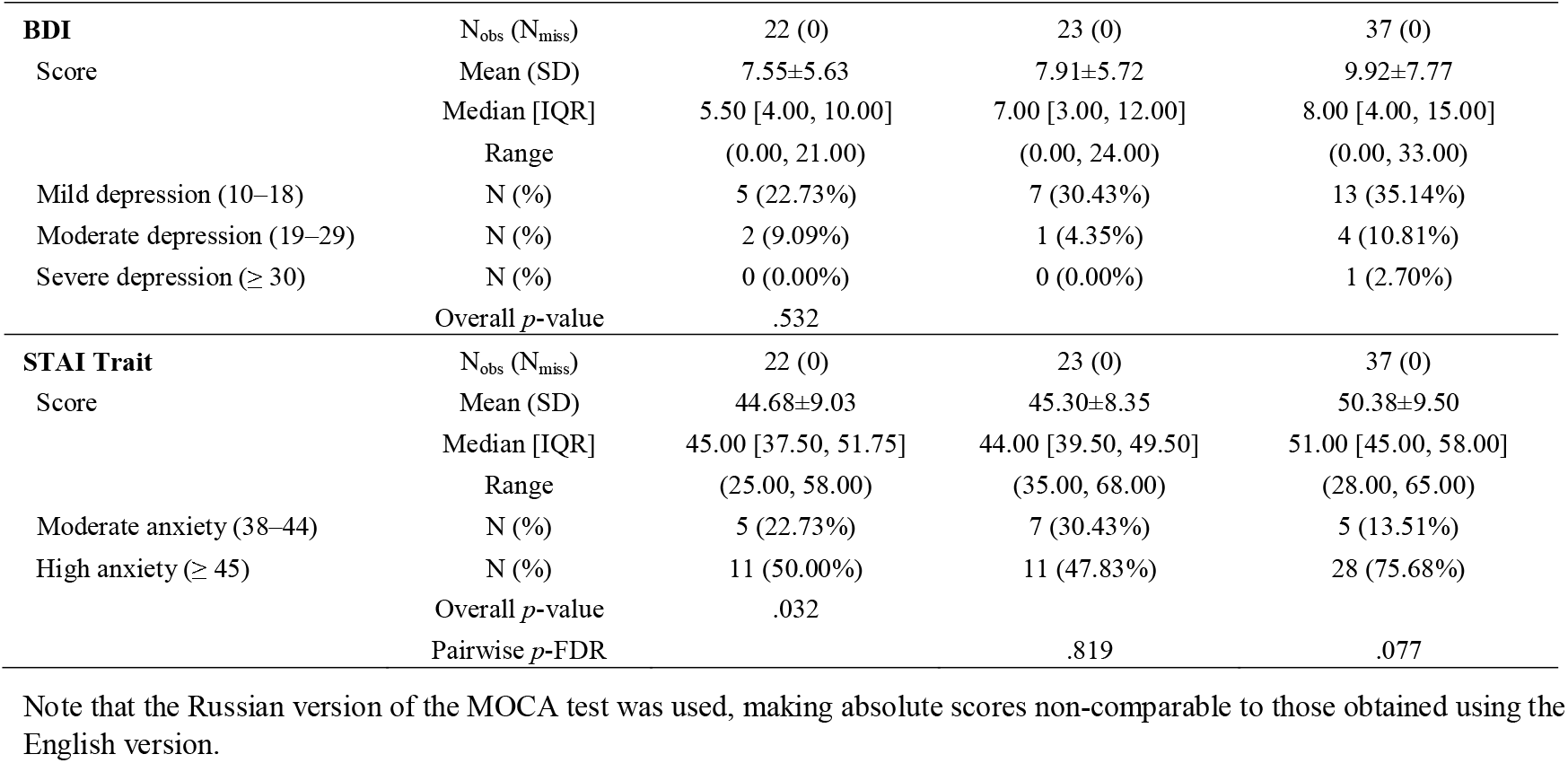
Characteristics of the Study Sample.

The two middle-aged groups were matched on general cognitive ability, depressive symptoms, arterial hypertension status, and body mass index (BMI).

### Altered Emotional Differentiation in Early cSVD

Groups differed in their ability to differentiate emotional experiences during the social emotion fMRI task. Middle-aged adults with cSVD showed lower emotional differentiation than those without cSVD (χ^2^ = 6.27, *p* =.044; *p*-FDR =.042 for SVD− vs. SVD+; Figure 1A), suggesting reduced emotional granularity. These experimental findings were mirrored in self-reported emotional insight. On TAS-20, which measures difficulties in identifying and describing emotions in daily life, the middle-aged SVD+ group scored higher than those without cSVD, while middle-aged SVD+ adults outperformed young adults (F = 5.64, *p* =.005; *p*-FDR =.003 and.046, respectively; Figure 1B).

**Figure 1.**
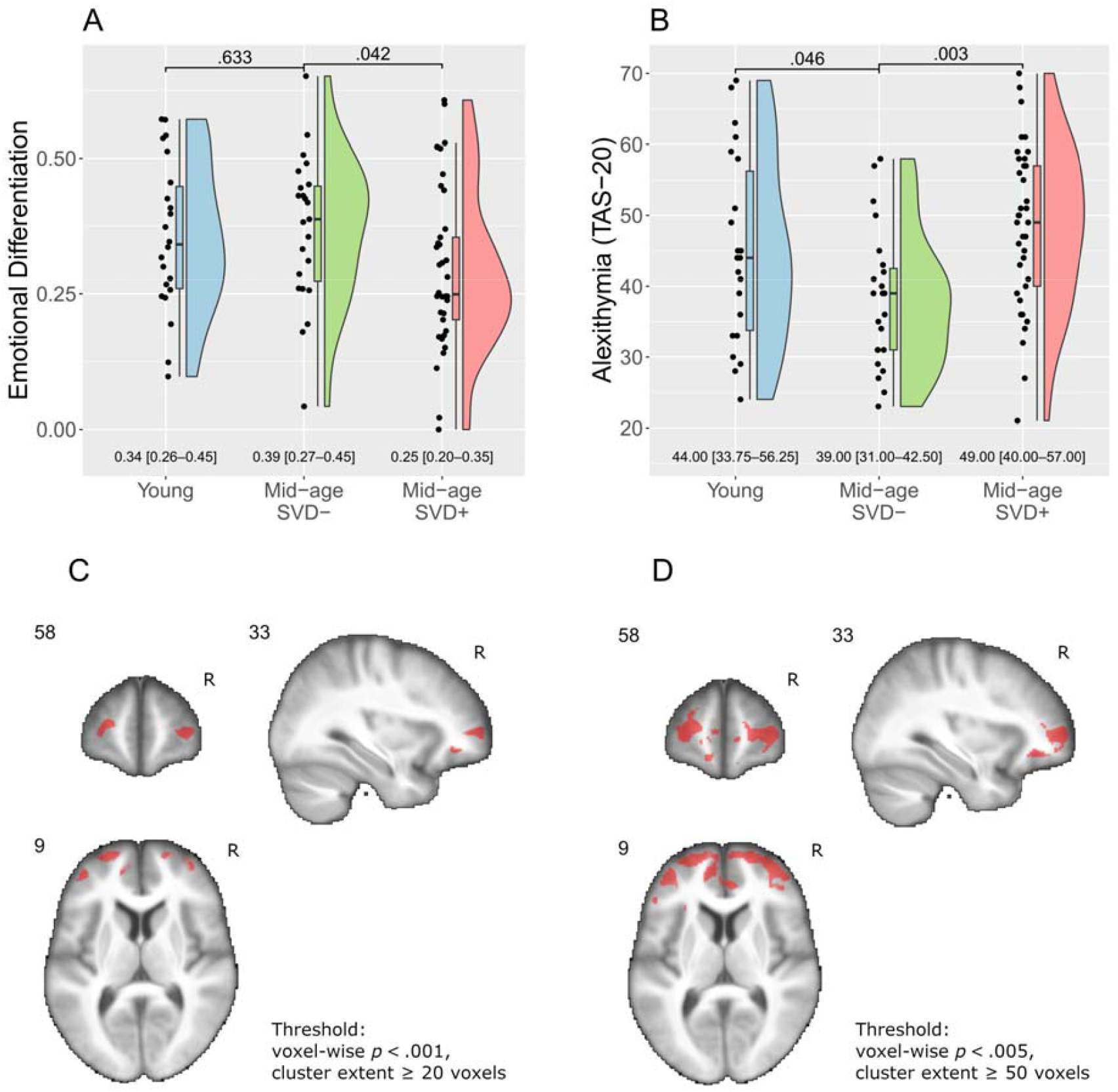
Alterations to Emotional Differentiation in Early Small Vessel Disease. Panels **A** and **B** indicate reduced emotional differentiation and heightened alexithymia (Toronto Alexithymia Scale, TAS-20) in early small vessel disease (middle-aged SVD+ vs. SVD−), but not healthy ageing (mid-age SVD− vs. young adults). On raincloud plots, dots correspond to individual observations; bar plots and numeric values represent median [interquartile range]; and density plots show data distribution. Significance is indicated for pairwise group comparisons, false discovery rate corrected. Panels **C** and **D** show the functional activation supporting emotional differentiation, overlaid over the mean group anatomy (Emotion vs. Pixel contrast; emotional differentiation as a second-level covariate; N= 82). Fragmented activation in the prefrontal cortex is seen at voxel-wise *p* <.001, with no cluster-level significance (C). At voxel-wise *p* <.005, the distributed effect across bilateral prefrontal cortex forms a cluster of 16,408 mm^3^, significant at the cluster-level *p*-FDR <.0001 (D).

To understand the neural basis of these differences in emotional differentiation, we examined the associated functional activation patterns (Emotion > Pixel contrast, second-level covariate analysis for emotional differentiation). At a conservative voxel-wise threshold of *p* <.001, several small clusters emerged in the prefrontal cortex, though none survived cluster-level correction (Figure 1C). When the threshold was lowered to *p* <.005, a distributed pattern of activation was observed across the bilateral prefrontal cortex (Figure 1D), forming an extended cluster of 2,051 voxels (16,408 mm^3^) at cluster-level *p*-FDR <.0001 with activation peaks in the right orbitofrontal cortex (30, 42, −8), right lateral prefrontal cortex (34, 60, 4), and left lateral prefrontal cortex (−22, 60, 12). This suggests that fine-grained emotional differentiation may be supported by a distributed prefrontal network.

### Loss of the Age-Related Positivity Effect in Early cSVD

Both middle-aged SVD+ and young adults labelled their emotional experiences more negatively than middle-aged SVD− adults. This was reflected in the mean valence ratings during the social emotion task, which differed significantly across groups (χ^2^ = 11.74, *p* =.003), with higher ratings in the SVD− group (*p*-FDR =.004 vs. young; *p*-FDR =.023 vs. SVD+; Figure 2A). Closer examination of the valence rating distributions (Figure 2B) revealed that young adults gave more negative and fewer positive ratings than SVD− middle-aged adults (*p*-FDR =.033 and.038, respectively), while SVD+ middle-aged adults gave more negative ratings than those without cSVD (*p*-FDR =.016), but did not differ in positive ratings (*p*-FDR =.788). Notably, only the middle-aged SVD− group showed a reliable shift toward more positive responses overall (*p* =.0002). Thus, the adaptive age-related positivity effect appeared disrupted by early cSVD.

**Figure 2.**
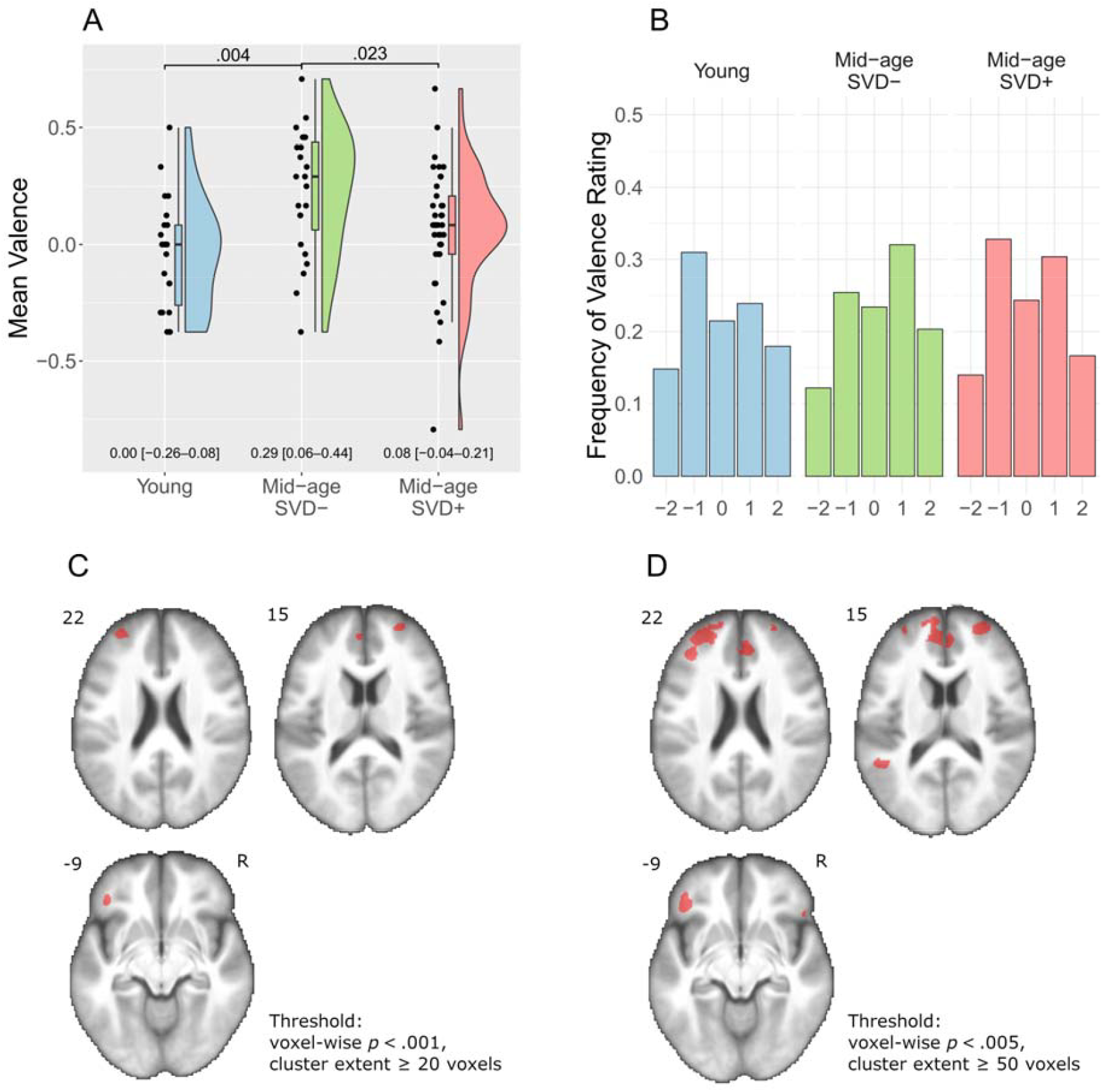
Alterations to the Positivity Effect in Early Small Vessel Disease. Panel **A** shows the age-related positivity effect in healthy ageing (mid-age SVD− vs. young adults) and its disruption by early SVD (middle-aged SVD+ vs. SVD−), as evidenced by the mean valence ratings. On the raincloud plot, dots correspond to individual observations; bar plots and numeric values represent median [interquartile range]; and density plots show data distribution. Significance is indicated for pairwise group comparisons, false discovery rate corrected. Panel **B** shows an altered distribution of valence ratings in early SVD, with a higher frequency of negative ratings compared to SVD− middle-aged adults. Panels **C** and **D** show the functional activation supporting positivity, overlaid over the mean group anatomy (Emotion vs. Pixel contrast; mean valence as a second-level covariate; N = 82). The fragmented effect in the prefrontal cortex did not reach the cluster-level significance.

Second-level covariate analysis (Emotion > Pixel contrast) did not identify significant clusters linked to mean valence ratings at the whole-brain level. Exploratory voxel-wise analyses revealed a fragmented and spatially diffuse pattern of activation across the prefrontal cortex, particularly at more liberal statistical thresholds (Figure 2C and D), suggesting that the neural correlates of emotional valence may be subtle and distributed or exhibit high inter-individual variability.

### Disrupted Neurovisceral Integration in Early cSVD

To assess neurovisceral integration, we examined how subjective arousal modulated BOLD responses in the right ventral anterior insula (first-level parametric modulation analysis; Figure 3A). Modulation strength differed across groups (Kruskal–Wallis χ^2^ = 6.19, *p* =.045; Figure 3B). Among middle-aged adults, those with cSVD showed weaker insular encoding of arousal (*p*-FDR =.019), suggesting disrupted neurovisceral integration. The difference between young and middle-aged SVD− adults was not significant (*p*-FDR =.164), indicating that ageing alone does not account for this effect. Importantly, individuals with stronger insular encoding of arousal also exhibited a more marked decrease in heart rate throughout the task (β = −0.006, *p* <.001; Table 2), linking intact neurovisceral integration to more effective physiological adaptation.

**Figure 3.**
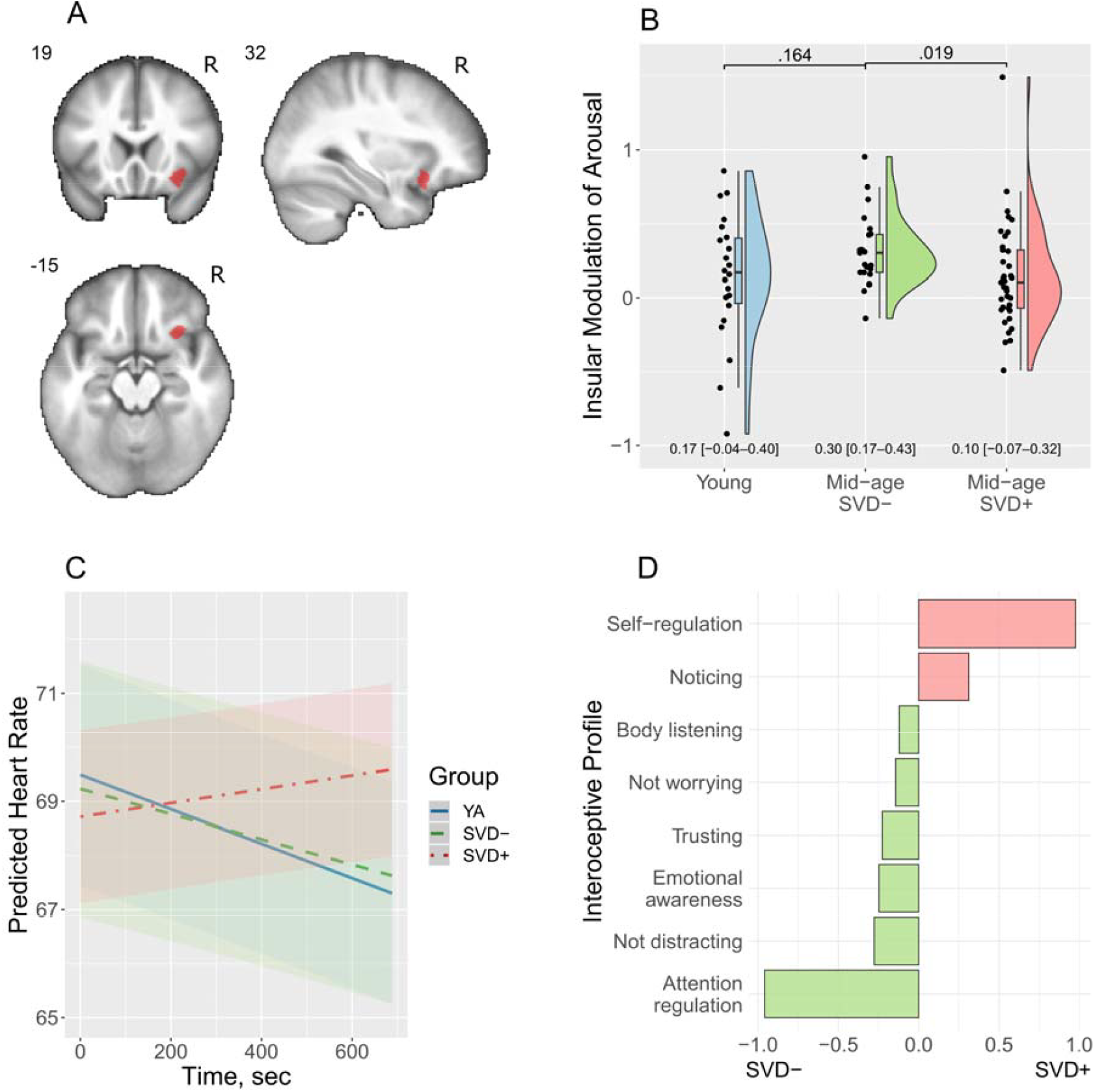
Alterations to Neurovisceral Integration in Early Small Vessel Disease. Panel **A** shows the region of interest (ROI) in the right ventral anterior insula that supports neurovisceral integration, selected for analysis. ROI data were estimated from first-level analyses using arousal as a parametric modulator. On panel **B**, reduced insular modulation of arousal is observed in early small vessel disease (SVD), but not in healthy ageing. On the raincloud plot, dots correspond to individual observations; bar plots and numeric values represent median [interquartile range]; and density plots show data distribution. Significance is indicated for pairwise group comparisons, false discovery rate corrected. Panel **C** shows altered heart rate dynamics during the social emotion task in early SVD, estimated from a linear mixed-effects model with a time-by-group interaction (shades indicate standard errors). In SVD+ participants, heart rate gradually increases throughout the task, while young adults (YA) and SVD− middle-aged adults show adaptive physiological habituation. On panel **D**, middle-aged adults with and without SVD demonstrate different interoceptive profiles, as indicated by the Multidimensional Assessment of Interoceptive Awareness (MAIA-2). Discriminant function scores are displayed on the x-axis, indicating how strongly MAIA-2 subscales align with the SVD− and SVD+ profiles.

**Table 2.**
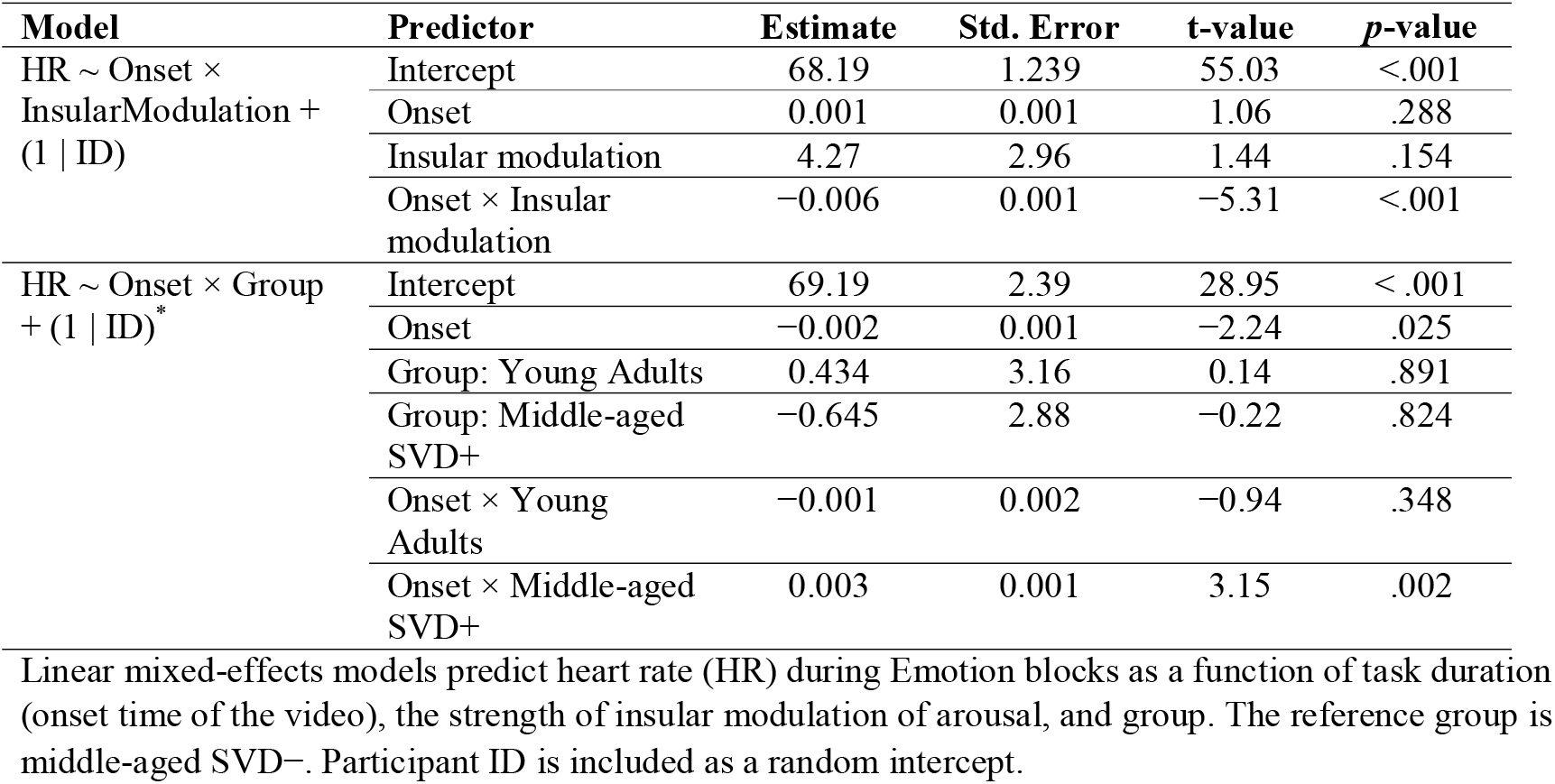
Heart Rate Dynamics are Shaped by Insular Modulation of Arousal and are Altered in Small Vessel Disease.

A linear mixed-effects model was used to examine how age and cSVD status influenced heart rate dynamics during the social emotion task (Emotion trials), including group, video onset time, and their interactions (Table 2). Across all participants, heart rate decreased over time (β =−0.002, *p* =.025), consistent with physiological adaptation. This dynamic was reversed in middle-aged adults with cSVD, who showed a significant positive time-by-group interaction (β = 0.003, *p* =.002), indicating an increase in heart rate over the course of the task and suggesting diminished habituation to emotional stimuli (Figure 3C).

To explore differences in interoceptive self-report profiles, we applied LDA and multivariate testing to MAIA-2 subscales. No significant multivariate differences were observed between young and middle-aged SVD− adults (Wilks’ λ = 0.749, *p* =.189), and classification accuracy was low (56%). In contrast, interoceptive profiles of middle-aged SVD+ adults differed from those without cSVD (Wilks’ λ = 0.707, *p* =.017), with a classification accuracy of 68%. According to the discriminant function, the most distinguishing features of cSVD were reduced *attention regulation* and *emotional awareness*, as well as elevated *self-regulation* and *noticing* (Figure 3D). These findings suggest that cSVD is associated with a shift in interoceptive experience that is not attributable to ageing alone and may reflect an emerging disruption in self-regulatory interoceptive processing.

### Differences Not Explained by Other Factors

To determine whether the observed group differences could be explained by other demographic, clinical, or cognitive variables, we conducted a series of exploratory analyses.

Gender was not associated with emotional differentiation, positivity, or neurovisceral integration (Table S4), suggesting that the gender imbalance between groups did not contribute to the findings. Arterial hypertension, similarly to cSVD, was linked to lower self-reported ability to regulate interoceptive attention (Table S5), but showed no significant associations with other measures of interest. Multivariate group differences in interoceptive profiles remained significant after adjusting for arterial hypertension (Wilks’ λ = 0.691, *p* =.014), which itself was no longer a significant predictor (Wilks’ λ =.756, *p* =.070). This suggests that the interoceptive alterations observed in cSVD are not explained by comorbid hypertension.

Depression and trait anxiety scores were correlated with alexithymia (Figure S4). However, post hoc models showed no significant main effects or interactions with group, indicating these affective symptoms did not account for the group differences. BMI was unrelated to the variables of interest, while grey matter volume showed a weak negative correlation with positivity, which did not remain in post hoc modelling.

In the cognitive examination, middle-aged SVD+ vs. SVD− adults scored lower in the Rey-Osterrieth Complex Figure Test (delayed recall), Verbal Fluency, and Trail Making tests (Table S6). Since verbal fluency was also associated with emotional differentiation, alexithymia and insular modulation of arousal (Figure S4), we tested whether it might explain the group effects. For both emotional differentiation and alexithymia, verbal fluency showed no significant main or interaction effects (p >.39). For insular modulation, an interaction between group and verbal fluency was observed (t = 2.14, p =.037). However, the group effect remained significant (t =−2.38, p =.021), suggesting that verbal fluency moderated, but did not mediate, the group difference.

## Discussion

The present study demonstrates disruption of neurovisceral integration and emotional processing in early cSVD, distinguishing it from healthy ageing. Middle-aged adults with cSVD showed reduced emotional differentiation, loss of the age-related positivity effect, impaired insula-mediated modulation of arousal, and lack of physiological habituation. These changes were evident prior to clinically significant cognitive decline, suggesting that affective alterations may represent an early marker of microvascular brain injury.

### Disrupted Emotional Processing and Neurovisceral Integration in Early cSVD

*Emotional differentiation*, operationalised as the granularity of affective ratings during the fMRI task, was preserved in healthy ageing but reduced in early cSVD (Figure 1). Consistently, participants with cSVD reported greater difficulty identifying and differentiating emotions in daily life. Emotional differentiation enables precise mapping between affective states and physiological responses, supporting efficient regulatory control.^40, 41^ Reduced differentiation in cSVD may therefore reflect impaired integration of cortical and subcortical networks involved in affective evaluation and response selection.^3^ In line with this interpretation, lower emotional differentiation was associated with reduced engagement of prefrontal regions during the social emotion fMRI task. Distributed bilateral activation across lateral and orbitofrontal prefrontal regions may reflect the diversity of neural representations underlying different top-down regulation strategies.^42^

Middle-aged adults without cSVD showed the typical *age-related positivity effect*, whereas this pattern was absent in those with early cerebrovascular injury (Figure 2). The group difference was driven by a higher proportion of negative ratings among adults with cSVD, suggesting reduced capacity to downregulate negative affect. Neurobiological studies link age-related positivity to medial prefrontal modulation of limbic activity, dependent on intact fronto-subcortical circuitry.^43, 44^ Its disruption in early cSVD is thus consistent with subtle network-level dysfunction affecting regulatory control mechanisms.

Linking altered emotional and physiological self-regulation, early cSVD was associated with disrupted *neurovisceral integration*, as evidenced by fMRI, heart rate dynamics, and self-reports of interoception (Figure 3). Adults with cSVD showed reduced modulation of arousal by the right ventral anterior insula. This region encodes arousal intensity and functions as a key node within the hierarchical network that integrates interoceptive information into representations of emotional states and enables differentiated self-regulation.^39^ Impaired insular functioning was associated with diminished psychophysiological adaptation during emotional processing. Participants with cSVD, unlike other groups, showed a progressive increase in heart rate throughout the social emotion task, suggesting impaired cardiovascular reactivity. Experimental findings were paralleled by changes in self-report interoceptive profiles. Middle-aged adults with cSVD showed a shift from embodied attention regulation and emotional awareness to more effortful, detached, top-down self-regulation strategies.

Overall, cSVD was associated with disruptions in the systems supporting neurovisceral integration and emotional processing across the neural, physiological, and behavioural levels. Within this regulatory dysfunction, impaired cardiovascular reactivity may represent a pathway to increased vascular burden, providing a potential mechanistic connection to white matter injury and cognitive decline.

### Independence from Confounders and Relevance to Everyday Functioning

Study findings persisted after correction for gender, cardiovascular risk factors such as hypertension and BMI, and cognitive performance (Tables S4, S5, and S6; Figure S4), suggesting an independent role for affective alterations in cSVD. Notably, verbal fluency moderated the relationship between cSVD and insular modulation of arousal. This finding may reflect the role of verbal mediation in bodily self-regulation^45^ and is consistent with our earlier observation of reduced resting-state connectivity between the salience and language networks in cSVD-related emotional dysregulation.^3^ The revealed role of verbal processing points to a trajectory in which advancing cognitive decline could further undermine neurovisceral integration.

The study aimed for ecological validity, a goal difficult to achieve in neuroimaging research. Firstly, we designed the fMRI task to approximate real-life social stressors in a multisensory, dynamic way. Rather than being stimulated by abstract, static threat cues, participants viewed videos of actors speaking to them in a pleasant or confrontational manner. This social emotion task aimed to mimic everyday interpersonal interactions. Secondly, we complemented the experimental paradigm with self-report questionnaires assessing the same processes as they occur in daily life. This allowed us to evidence the real-world relevance of alterations in emotional processing and neurovisceral integration.

### Limitations and Future Directions

Due to the cross-sectional nature of the study, it remains unclear whether the observed differences between middle-aged adults with and without cSVD were secondary to emerging brain injury, reflected a trait vulnerability that predisposed to cSVD, or a combination of both. In a previous longitudinal study, nocturnal heart rate variability was predictive of four-year cSVD progression, suggesting a mechanistic role for disrupted neurovisceral integration, along with evidence from general cardiovascular research.^7, 46^ Further longitudinal studies are required to clarify the causal relationships. Research should also aim for more demographically balanced and ethnically diverse samples to enhance generalisability.

## Conclusions

This study advances understanding of age-related vascular brain injury and its underexplored affective aspects. Experimental findings from an ecologically grounded fMRI task, paralleled by self-reports of daily functioning, indicate pervasive alterations across behavioural, neural, and physiological levels. Disruption of emotional differentiation, positivity, and neurovisceral integration may represent candidate biomarkers of emerging cSVD and mechanistic pathways to cognitive decline. These alterations also highlight targets for behavioural and neurophysiological interventions to preserve brain health across the lifespan, delineating a translational avenue for early intervention.

## Author Contribution Statement

Conceptualisation: OD, LD, GA, EN, EK, MK. Data curation: EN, VK, DA. Formal analysis: OD. Funding acquisition: LD, OD. Investigation: VT, EN, VK, DA. Methodology: OD, LD, GA, VK, DA, EK, MK. Project administration: OD, GA, LD. Visualisation: OD. Writing – original draft: OD, LD, GA. Writing – review & editing: All authors.

## Statements and Declarations

### Ethical considerations

The study protocol was approved by the Ethics Committee and the Institutional Review Board of the Russian Center of Neurology and Neurosciences (protocols 4-6/19, 1-3/20, and 4-3/21).

### Consent to participate

All participants gave written informed consent to participate in the study.

### Consent for publication

Not applicable

### Data availability

The data supporting this study are available from the corresponding author upon reasonable request.

