## Supplementary Information for "Disrupted Neurovisceral Integration and Emotional Processing in Early Cerebral Small Vessel Disease"

Olga R. Dobrushina[a, b], Larisa A. Dobrynina[a], Galina A. Arina[c], Evgenia S. Novikova[c], Mariia M. Tsypushtanova[c], Angelina G. Makarova[c], Mariia V. Gubanova[c], Viktoriya V. Trubitsyna[c], Vlada V. Aristova[a, c], Daria A. Kazantseva[a, c], Elena I. Kremneva[a, c], Marina V. Krotenkova[a, c]

[a] Russian Center of Neurology and Neurosciences, 80 Volokolamskoe, Moscow, Russia, 125367
[b] University of Strathclyde, 40 George Street, Glasgow, UK, G1 1QE
[c] M.V. Lomonosov Moscow State University, Faculty of Psychology, 11 Mochovaya, Moscow, Russia, 125009

Correspondence:
Olga Dobrushina, 40 George Street, Glasgow, UK, G1 1QE,

### Evaluation of the Social Emotion Task

This supplementary section provides an evaluation of the novel social emotion task to assess its performance and construct validity, performed across the entire sample of 82 participants.

#### *Arousal and Valence Ratings*

*Arousal* and *valence* ratings reflected the expected pattern of responses to actors' emotional expressions (see Table S1). We used linear mixed-effects models with participant as a random effect to test whether ratings varied by displayed emotion. Both arousal and valence ratings showed significant main effects of emotion ($p < .0001$). Joy and anger evoked the highest arousal, while irritation and pleasure were associated with lower arousal. Post hoc pairwise comparisons (Tukey-adjusted) indicated that all differences were significant ($p < .0001$) except between anger and joy. Valence ratings also aligned with expected emotional polarity: joy and pleasure were rated as most positive, irritation as near neutral, and anger as negative. All pairwise comparisons were significant ($p < .0001$), except between joy and pleasure ($p = .08$). The observed dissociation between arousal and valence ratings across emotion categories supports discriminant validity of the rating responses.

A separate mixed-effects model confirmed a small but statistically significant positive association between valence and arousal ($\beta = 0.065$, $p = .003$), indicating that more positively perceived stimuli were associated with slightly higher arousal. *Emotional differentiation*, quantified as the variance in absolute arousal and valence ratings across trials, was negatively correlated with alexithymia (Spearman's $\rho = -0.28$, $p = .010$), supporting the construct validity[1].

**Table S1. Arousal and Valence Ratings by Displayed Emotion**

| Emotion | Statistics | Arousal rating | Valence rating |
|---|---|---|---|
| **Pleasure** | Mean±SD | −0.01±1.17 | 0.82±0.94 |
| | Median [IQR] | 0.00 [−1.00, 1.00] | 1.00 [0.00, 1.00] |
| **Joy** | Mean±SD | 1.03±1.08 | 0.96±1.10 |
| | Median [IQR] | 1.00 [1.00, 2.00] | 1.00 [0.00, 2.00] |
| **Irritation** | Mean±SD | −0.49±1.12 | −0.29±0.90 |
| | Median [IQR] | −1.00 [−1.00, 0.00] | 0.00 [−1.00, 0.00] |
| **Anger** | Mean±SD | 0.95±1.11 | −1.15±0.80 |
| | Median [IQR] | 1.00 [1.00, 2.00] | −1.00 [−2.00, −1.00] |

#### *Heart Rate Dynamics*

To assess physiological engagement during the social emotion task, we examined heart rate dynamics. A linear mixed-effects model tested the effects of subjective arousal and time (video onset) on heart rate. The model revealed a significant overall decline in heart rate throughout the task ($\beta = -0.0011$, $p = .047$), consistent with task-related autonomic adjustment. Additionally, a significant interaction between arousal and onset ($\beta = 0.0009$, $p = .030$)

indicated that trials rated as more arousing were associated with a relative attenuation of the heart rate decrease over time. This suggests that subjective emotional arousal was reflected in autonomic responses during the task. The addition of valence to the model did not result in improved performance. These results support the task's capacity to evoke and capture individual differences in emotional reactivity at the physiological level.

*Functional Activation*

The *Emotion* > *Pixel* contrast revealed bilateral activation in temporal, frontal, and occipital regions involved in the perception of multimodal social stimuli, as well as emotional awareness and regulation (see Table S2 and Figure S2), consistent with previous findings on socio-emotional perception[2,3]. *Arousal modulation* was associated with activation in the insula, occipital cortex, and superior parietal lobule, consistent with heightened perceptual and interoceptive processing during emotionally arousing trials (see Table S3 and Figure S3). These regions have been previously implicated in salience detection[4], visuo-emotional integration[5], and physiological arousal monitoring[6,7]. No significant activation was associated with *valence modulation*.

*Conclusion*

Overall, behavioural, physiological, and neural responses support the construct validity of the social emotion task. Ratings followed theoretically expected patterns and differentiated between emotion categories; arousal responses correlated with heart rate and were reflected in activation of salience and interoceptive regions. The task captures individual differences in emotional reactivity across multiple modalities and offers higher ecological validity by evaluating participants' emotional responses to dynamic, multimodal social cues, mimicking everyday interpersonal interactions.

**Table S2. Activation Clusters for the Emotion > Pixel Contrast**

| Region | Peak MNI coordinates (x, y, z) | Cluster Size, voxels (2×2×2 mm3) | Cluster-level *p*-FDR |
|---|---|---|---|
| **Right superior temporal gyrus, middle and inferior occipital gyri** | 62 −20 2 | 5707 | < .001 |
| **Left superior temporal gyrus, middle and inferior occipital gyri** | −60 −24 4 | 5644 | < .001 |
| **Left inferior temporal gyrus** | −44 −48 −20 | 226 | .007 |
| **Right frontal operculum** | 54 28 2 | 430 | .001 |
| **Right precentral gyrus** | 54 0 48 | 299 | .003 |
| **Medial prefrontal cortex** | 6 60 26 | 999 | < .001 |
| **Left precentral gyrus** | −52 −4 48 | 210 | .009 |
| **Left frontal operculum** | −40 26 −12 | 525 | < .001 |

*Voxel-level threshold was set at p < .001, with clusters considered significant at p-FDR < .05.*

**Table S3. Activation Clusters for Arousal Modulation**

| Region | Peak MNI coordinates (x, y, z) | Cluster Size, voxels (2×2×2 mm3) | Cluster-level *p*-FDR |
|---|---|---|---|
| **Left insula, left superior temporal sulcus, left middle temporal gyrus** | −50 26 0 | 3910 | < .001 |
| **Right occipital pole** | 16 −86 −2 | 292 | < .001 |
| **Left lingual gyrus** | −16 −48 −4 | 624 | < .001 |
| **Left occipital pole** | −28 −92 −6 | 288 | < .001 |
| **Left superior parietal lobule** | −30 −64 60 | 274 | .001 |
| **Right anterior insula** | 30 20 −14 | 278 | < .001 |
| **Left precentral** | −42 −24 48 | 425 | < .001 |

*Voxel-level threshold was set at p < .001, with clusters considered significant at p-FDR < .05.*

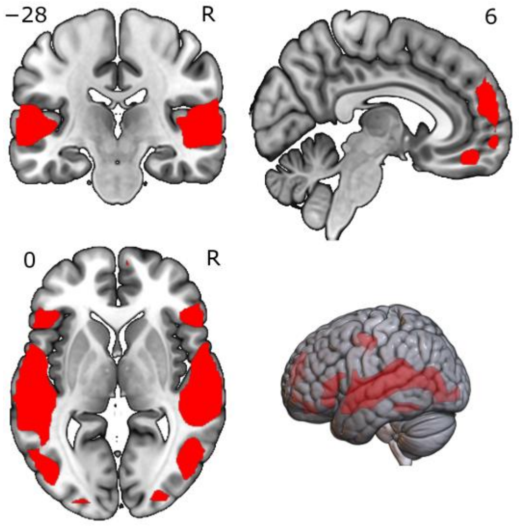


**Figure S2. Activation map for the Emotion > Pixel contrast**

*Voxel-level threshold: p < .001; cluster-level p-FDR < .05.*

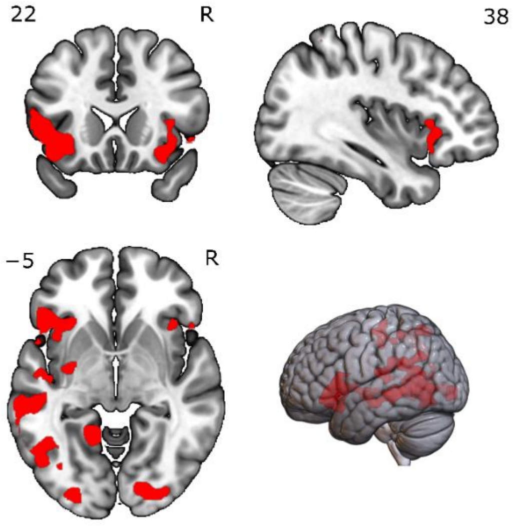


**Figure S3. Activation map for arousal modulation**

*Voxel-level threshold: p < .001; cluster-level p-FDR < .05.*
